# mangal - making ecological network analysis simple

**DOI:** 10.1101/002634

**Authors:** Timothée Poisot, Benjamin Baiser, Jennifer A. Dunne, Sonia Kéfi, François Massol, Nicolas Mouquet, Tamara N. Romanuk, Daniel B. Stouffer, Spencer A. Wood, Dominique Gravel

## Abstract

The study of ecological networks is severely limited by (i) the difficulty to access data, (ii) the lack of a standardized way to link meta-data with interactions, and (iii) the disparity of formats in which ecological networks themselves are represented. To overcome these limitations, we conceived a data specification for ecological networks. We implemented a database respecting this standard, and released a R package (`rmangal`) allowing users to programmatically access, curate, and deposit data on ecological interactions. In this article, we show how these tools, in conjunctions with other frameworks for the programmatic manipulation of open ecological data, streamlines the analysis process, and improves eplicability and reproducibility of ecological networks studies.

## Introduction

Ecological networks are efficient representations of the complexity of natural communities, and help discover mechanisms contributing to their persistence, stability, resilience, and functioning. Most of the early studies of ecological networks were focused on understanding how the structure of interactions within one location affected the ecological properties of this local community. They revealed the contribution of average network properties, such as the buffering impact of modularity on species loss (Pimm et al. 1991,???), the increase in robustness to extinctions along with increases in connectance (Dunne et al. 2002), and the fact that organization of interactions maximizes biodiversity (Bastolla et al. 2009). New studies introduced the idea that networks can vary from one locality to another. They can be meaningfully compared, either to understand the importance of environmental gradients on the presence of ecological interactions (Tylianakis et al. 2007), or to understand the mechanisms behind variation itself (Poisot et al. 2012, 2014). Yet, metaanalyses of numerous ecological networks are still extremely rare, and most of the studies comparing several networks do so within the limit of particular systems (Schleuning et al. 2011, Dalsgaard et al. 2013, Poisot et al. 2013, Chamberlain et al. 2014, Olito and Fox 2014). The severe shortage of publicly shared data in the field also restricts the scope of large-scale analyses.

It is possible to predict the structure of ecological networks, either using latent variables (Rohr et al. 2010, Eklöf et al. 2013) or actual trait values (Gravel et al. 2013). The calibration of these approaches require accessible data, not only about the interactions, but about the traits of the species involved. Comparing the efficiency of different methods is also facilitated if there is a homogeneous way of representing ecological interactions, and the associated metadata. In this paper, we (i) establish the need of a data specification serving as a common language among network ecologists, (ii) describe this data specification, and (iii) describe `rmangal`, a `R` package and companion database relying on this data specification. The `rmangal` package allows to easily deposit and retrieve data about ecological interactions and networks in a publicly accessible database. We provide use cases showing how this new approach makes complex analyzes simpler, and allows for the integration of new tools to manipulate biodiversity resources.

## Networks need a data specification

Ecological networks are (often) stored as an *adjacency matrix* (or as the quantitative link matrix), that is a series of 0s and 1s indicating, respectively, the absence and presence of an interaction. This format is extremely convenient for *use* (as most network analysis packages, *e.g.* `bipartite`, `betalink`, `foodweb`, require data to be presented this way), but is extremely inefficient at *storing* meta-data. In most cases, an adjacency matrix provides information about the identity of species (in the cases where rows and columns headers are present) and the presence or absence of interactions. If other data about the environment (*e.g.* where the network was sampled) or the species (*e.g.* the population size, trait distribution, or other observations) are available, they are often either given in other files or as accompanying text. In both cases, making a programmatic link between interaction data and relevant meta-data is difficult and, more importantly, error-prone.

By contrast, a data specification (*i.e.* a set of precise instructions detailing how each object should be represented) provides a common language for network ecologists to interact, and ensures that, regardless of their source, data can be used in a shared workflow. Most importantly, a data specification describes how data are *exchanged*. Each group retains the ability to store the data in the format that is most convenient for in-house use, and only needs to provide export options (*e.g.* through an API, *i.e.* a programmatic interface running on a web server, returning data in response to queries in a pre-determined language) respecting the data specification. This approach ensures that *all* data can be used in metaanalyses, and increases the impact of data (Piwowar and Vision 2013). Data archival also offers additional advantages for ecology. The aggregation of local observations can reveal large-scale phenomena (Reichman et al. 2011), which would be unattainable in the absence of a collaborative effort. Data archival in databases also prevents data rot and data loss (Vines et al. 2014), thus ensuring that data on interaction networks – which are typically hard and costly to produce - continue to be available and usable.

## Elements of the data specification

The data specification introduced here (Fig. 1) is built around the idea that (ecological) networks are collections of relationships between ecological objects, and each element has particular meta-data associated with it. In this section, we detail the way networks are represented in the `mangal` specification. An interactive webpage with the elements of the data specification can be found online at http://mangal.io/doc/spec/. The data specification is available either at the API root (*e.g.* http://mangal.io/api/v1/?format=json), or can be viewed using the `whatls` function from the `rmangal` package. Rather than giving an exhaustive list of the data specification (which is available online at the aforementioned URL), this section serves as an overview of each element, and how they interact.

**Fig. 1:**
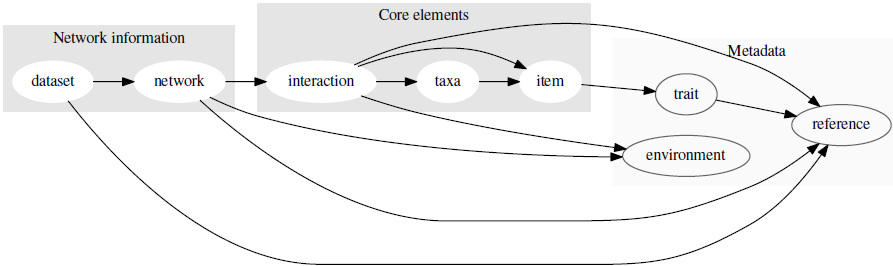
An overview of the data specification, and the hierarchy between objects. Every box corresponds to a level of the data specification. Grey boxes are nodes, blue boxes are interactions and networks, and green boxes are metadata. The **bold** boxes (`dataset, network, interaction, taxa`) are the minimal elements needed to brepresent a network.

We propose `JSON`, a user-friendly format equivalent to `XML`, as an efficient way to standardise data representation for two main reasons. First, it has emerged as a *de facto* standard for web platform serving data, and accepting data from users. Second, it allows strict *validation* of the data: a JSON file can be matched against a scheme, and one can verify that it is correctly formatted (this includes the possibility that not all fields are filled, as will depend on available data). Finally, `JSON` objects are easily and cheaply (memory-wise) parsed in the most commonly-used programming languages, notably `R` (equivalent to `list`) and `python` (equivalent to `dict`). For most users, the format in which data are transmitted is unimportant, as the interaction happens within `R` – as such, knowing how `JSON` objects are organized is only useful for those who want to interact with the API directly. As such, the `rmangal` package takes care of converting the data into the correct `JSON` format to upload them in the database.

Functions in the `rmangal` package are names after elements of the data specification, in the following way: `verb + Element`. `verb` can be one of `list`, `get`, or `patch`; for example, the function to get a particular network is `getNetwork`. The function to modify (patch) a taxon is `patchTaxa`. All of these functions return a *list* object, which means that chaining them together using, *e.g.* the `plyr` package, is time-efficient. There are examples of this in the use-cases.

## Node information

### Taxa

Taxa are a taxonomic entity of any level, identified by their name, vernacular name, and their identifiers in a variety of taxonomic services. Associating the identifiers of each taxa allows using the new generation of open data tools, such as `taxize` (Chamberlain and Szöcs 2013), in addition to protecting the database against taxonomic revisions. The data specification currently has fields for `ncbi` (National Center for Biotechnology Information), `gbif` (Global Biodiversity Information Facility), `tsn` (Taxonomic Serial Number, used by the Integrated Taxonomic Information System), `eol` (Encyclopedia of Life) and `bold` (Barcode of Life) identifiers. We also provide the taxonomic status, *i.e*. whether the taxon is a true taxonomic entity, a “trophic species”, or a morphospecies. Taxonomic identifiers can either be added by the contributors, or will be automatically retrieved during the automated curation routine.

### Item

An `item` is any measured instance of a taxon. Items have a `level` argument, which can be either `individual` or `population`; this allows representing both individual-level networks (*i.e.* there are as many `items` of a given taxa as there were individuals of this taxon sampled), and population-level networks. When `item` represents a population, it is possible to give a measure of the size of this population. The notion of `item` is particularly useful for time-replicated designs: each observation of a population at a time-point is an `item` with associated `trait` values, and possibly population size.

## Network information

All objects described in this sub-section can have a spatial position, information on the date of sampling, and references to both papers and datasets.

### Interaction

An `interaction` links two `taxa` objects (but can also link pairs of `items`). The most important attributes of `interactions` are the type of interaction (of which we provide a list of possible values, see *Supp. Mat.* 1), and its `ob_type`, *i.e.* how it was observed. This field helps differentiate direct observations, text mining, and inference. Note that the `obs_type` field can also take `confirmed absence` as a value; this is useful for, *e.g.*, “cafeteria” experiments in which there is high confidence that the interaction did not happen.

### Network

A `network` is a series of `interaction` objects, along with (i) information on its spatial position (provided at the latitude and longitude), (ii) the date of sampling, and (iii) references to measures of environmental conditions.

### Dataset

A dataset is a collection of one or several network(s). Datasets also have a field for data and papers, both of which are references to bibliographic or web resources that describe, respectively, the source of the data and the papers in which these data have been studied. Datasets or networks are the preferred entry point into the resources, although in some cases it can be meaningful to get a list of interactions only.

## Meta-Data

### Trait value

Objects of type `item` can have associated `trait` values. These consist in the description of the trait being measured, the value, and the units in which the measure was taken. As traits may have been measured at a different time and/or location that the interaction was, they have fields for time, latitude and longitude, and references to original publication and original datasets.

### Environmental condition

Environmental conditions are associated to datasets, networks, and interactions objects, to allow for both macro and micro environmental conditions. These are defined by the environmental property measured, its value, and the units. As traits, they have fields for time, latitude and longitude, and references to original publication and original datasets.

## References

References are associated to datasets. They accommodate the DOI, `JSON` or PubMed identifiers, or a URL. When possible, the DOI is preferred as it offers more potential to interact with other online tools, such as the *CrossRef* API.

## Use cases

In this section, we present use cases using the `rmangal` package for R, to interact with a database implementing this data specification, and serving data through an API (http://mangal.io/api/v1/). It is possible for users to deposit data into this database through the R package. Note that data are made available under a *CC*–0 *Waiver* (???). Detailed information about how to upload data are given in the vignettes and manual of the `rmangal` package. In addition, the `rmangal` package comes with vignettes explaining how users can upload their data into the database through `R`.

The data we use for this example come from Ricciardi et al. (2010). These data were previously available on the *Interaction Web DataBase* as a single xls file. We uploaded them in the `mangal` database at http://mangal.io/api/v1/dataset/2. The `rmangal` package can be installed this way:

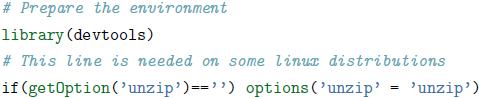

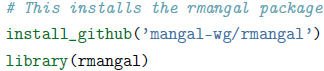

Once `rmangal` is installed and loaded, users can establish a connection to the database this way:

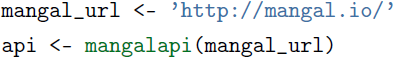

## Create taxa and add an interaction

In the first use-case, we will create an interaction between two taxa. We ask of readers *not* to execute this code as it is, but rather to use it as a template for their own analyses. A complete, step-by-step guide to upload is given in the vignettes of the `rmangal` package. Uploading anything requires an username and API key, which can be obtained at the following URL:http:/mangal.io/dashboard/login. Your API key be generated automatically after registration. You can use it to connect to the database securely:

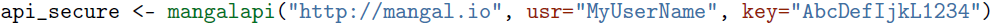

The first step is to create two taxa objects, with the species that we observed interacting:

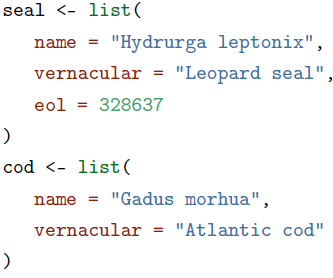

Now, we will send these two objects in the remote database:

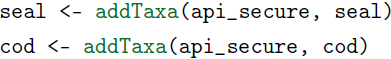

Note that it is suggested to overwrite the local copy of the object, because the database will *always* send back the remote copy. This makes the syntax of further addition considerably easier, as we show below. Once the two objects are created, we can create an interaction between them:

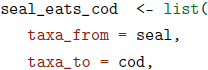

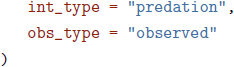

Then using the same approach, we can send this information in the remote database:

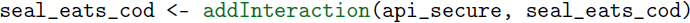

To create networks, datasets, etc, one needs follow the same procedure, as is explained in the online guide for data contributors, available at http://mangal.io/doc/upload/.

## Link-species relationships

In the first example, we visualize the relationship between the number of species and the number of interactions, which Martinez (1992) proposed to be linear (in food webs).

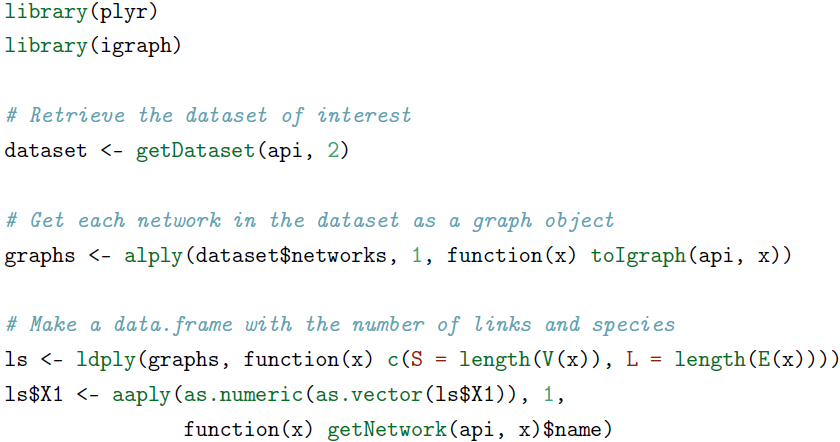

\## Error in eval(expr, envir, enclos): client error: (404) Not Found

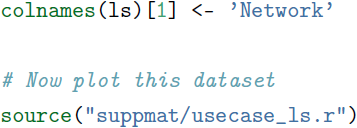

Getting the data to produce this figure requires less than 10 lines of code. The only information needed is the identifier of the network or dataset, which we suggest should be reported in publications as: “These data were deposited in the `mangal` format at `<URL>/api/v1/dataset/<ID>”` (where `<URL>` and `<ID>` are replaced by the corresponding values), preferably in the methods, possibly in the acknowledgements. To encourage data sharing and its recognition, we encourage users of the database to always cite the original datasets or publications.

**Fig. 2:**
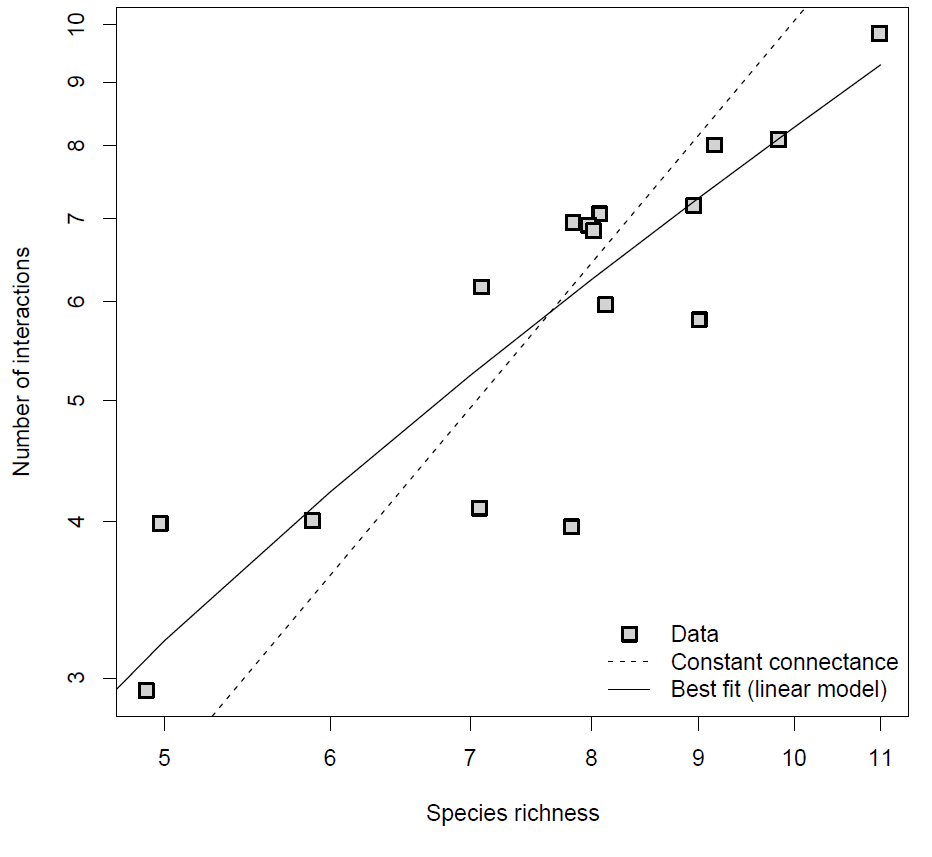
Relationship between the number of species and number of interactions in the anemonefish-fish dataset. Constant connectance refers to the hypothesis that there is a quadratic relationship between these two quantities.

## Network beta-diversity

In the second example, we use the framework of network *β*–diversity (Poisot et al. 2012) to measure the extent to which networks that are far apart in space have different interactions. Each network in the dataset has a latitude and longitude, meaning that it is possible to measure the geographic distance between two networks. For each pair of networks, we measure the geographic distance (in km), the species dissimilarity (*β*_*s*_), the network dissimilarity when all species are present (*β*_*WN*_), and finally, the network dissimilarity when only shared species are considered (*β*_*OS*_).

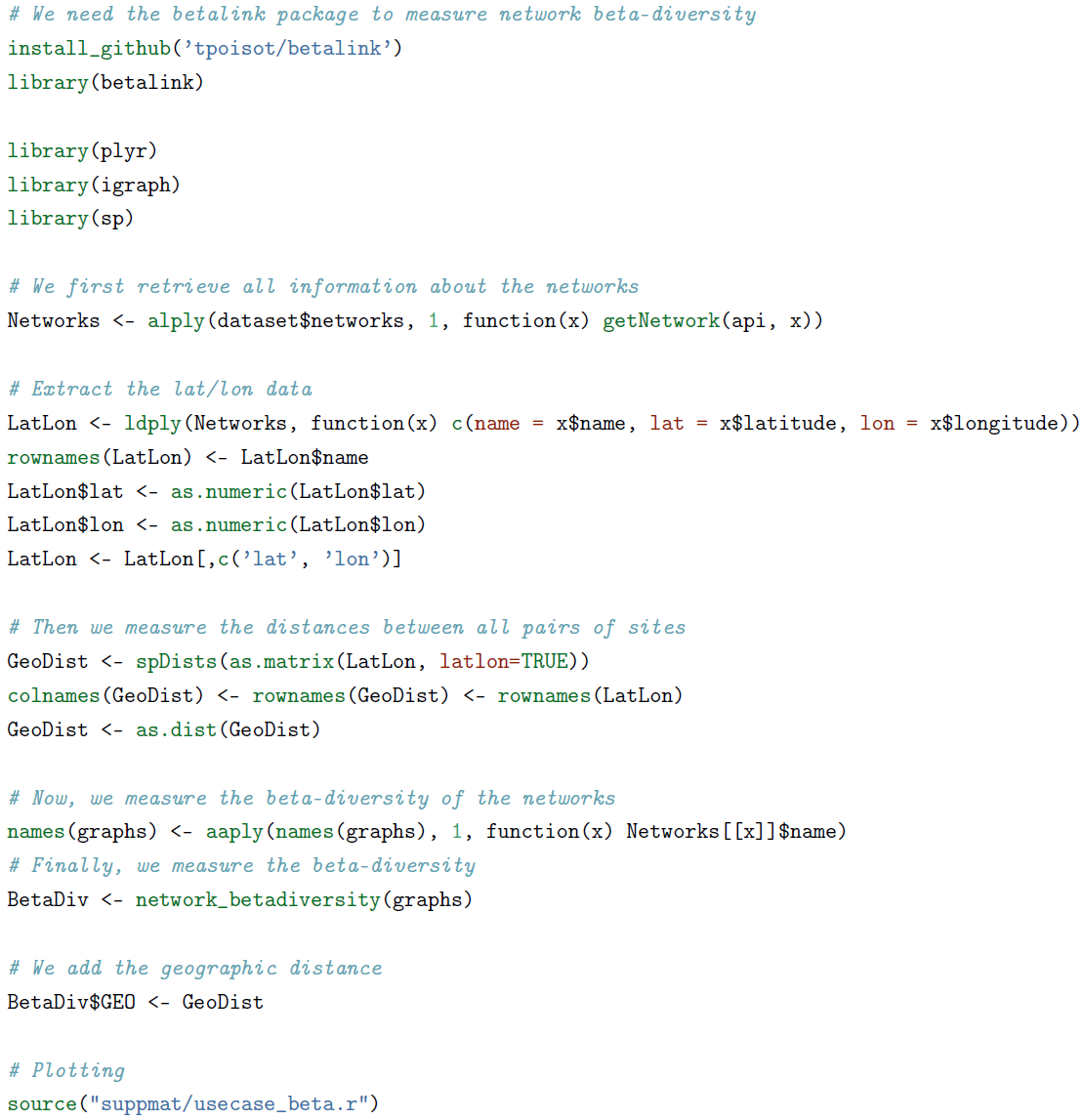

As shown in Fig. 3 while species dissimilarity and overall network dissimilarity increase when two networks are far apart, this is not the case for the way common species interact. This suggests that in this system, network dissimilarity over space is primarily driven by species turnover. The ease to gather both raw interaction data and associated meta-data make conducting this analysis extremely straightforward.

**Fig. 3:**
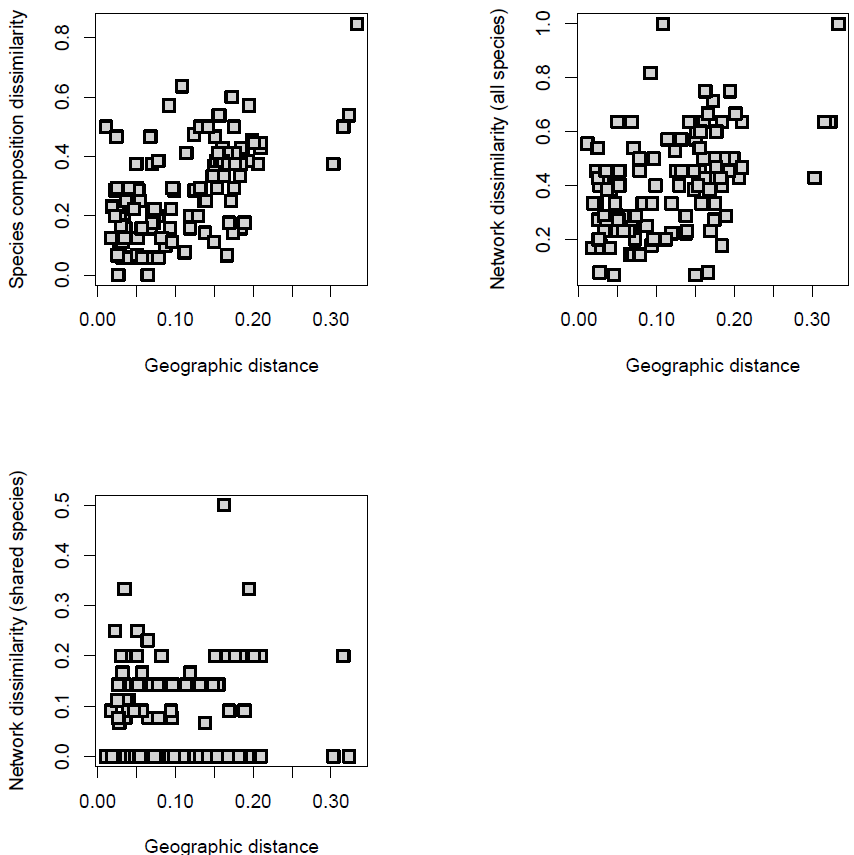
Relationships between the geographic distance between two sites, and the species dissimilarity, network dissimilarity with all species, and network dissimilarity with only shared species.

## Spatial visualization of networks

Bascompte (2009) uses an interesting visualization for spatial networks, in which each species is laid out on a map at the center of mass of its distribution; interactions are then drawn between species to show how species distribution determines biotic interactions. In this final use case, we propose to reproduce a similar figure (Fig. 4).

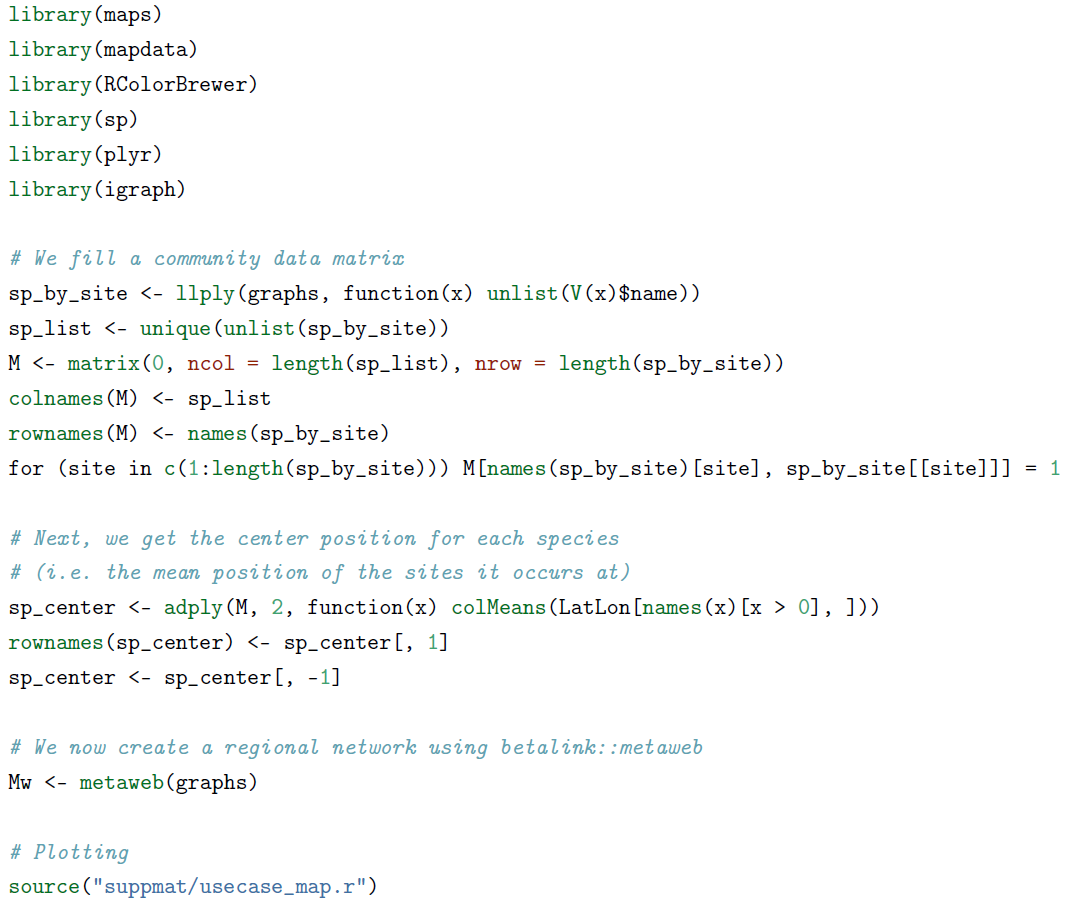

## Conclusions

The `mangal` data format will allow researchers to put together dataset with species interactions and rich meta-data, that are needed to address emerging questions about the structure of ecological networks. We deployed an online database with an associated API, relying on this data specification. Finally, we introduced `rmangal`, an R package designed to interact with APIs using the `mangal` format. We expect that the data specification will evolve based on the needs and feedback of the community. At the moment, users are welcome to propose such changes on the project issue page: https://github.com/mangal-wg/mangal-schemes/issues. A `python` wrapper for the API is also available at http://github.com/mangal-wg/pymangal/. Additionally, there are plans to integrate this database with *GLOBI,* so that data can be accessed from multiple sources (Poelen et al. 2014).

**Fig. 4:**
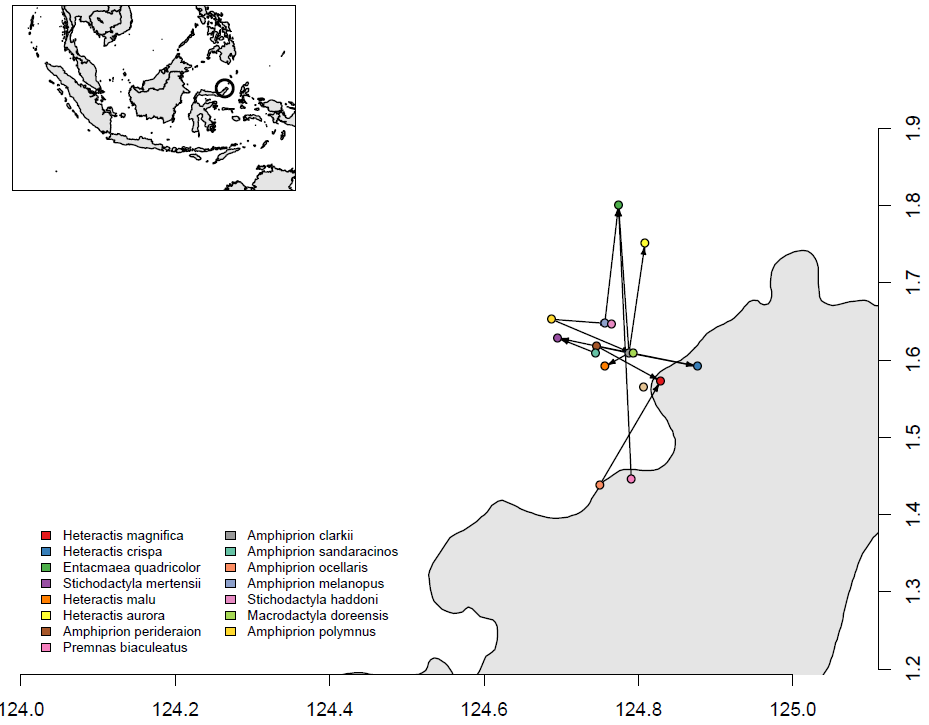
Spatial plot of a network, using the maps and `rmangal` packages. The circles in the inset map show the location of the sites. Each dot in the main map represents a species, with symbiotic mutualisms drawn between them. The land is in grey

## Acknowledgements

This paper was developed during a workshop hosted at the *Santa Fe Institute*. TP, DBS, and DG acknowledge funding from the Canadian Institute of Ecology and Evolution. We thank Scott Chamberlain and one anonymous reviewer for comments on the manuscript. TP is funded by a start-up grant from the Universite de Montreal. We thank the `rOpenSci` team and developers for inspiration.

